# Comprehensive analysis identifies histone 2A ubiquitination as a marker for aging and pan-cancer prognosis

**DOI:** 10.1101/2022.07.13.499990

**Authors:** Fen Zhang, Zhan Wang, Jiahang Zhang, Menghao Zhou, Yongxia Chen, Sainan Zhang, Zeyu Sun, Feiyang Ji

## Abstract

Ubiquitination is a post-translational modification (PTM) that induces protein degradation or function alteration and plays crucial roles in aging and cancer. Previous ubiquitinomes of aging mainly focused on how ubiquitination changes in drosophila and other lower animals, but how ubiquitination changes during the aging of higher animals and what causes these changes remain unclear. Here, we profiled whole-life ubiquitinome data of mouse brain, heart, liver, muscle, and spleen, and integratively analyzed the ubiquitinome data with RNA sequencing data. The results showed that the ubiquitination of protein, especially histone 2A (H2A), changed intensely during aging due to the regulated expression of E3 ligases (E3s) and deubiquitylating enzymes (DUBs). Then we developed two distinct H2A’ E3s/DUBs expression subtypes with different prognosis, DNA damage response (DDR), and tumor microenvironment cell infiltration degrees based on an unsupervised method in pan-cancer. In conclusion, our study provided temporal resolution ubiquitinome data of mouse aging and revealed the vital role of H2A ubiquitination in aging and tumor progression.

## Background

Ubiquitination is a post-translational modification (PTM) of proteins, which plays an obligate role in a multitude of physiological and pathological processes. The core molecule involved is a 76 amino-acid protein called ubiquitin, which can be linked to each other through 7 lysine residues and N-terminal methionine to form a polyubiquitin chain. The homeostasis of protein ubiquitination is maintained by the two hostile systems: ubiquitination and deubiquitination. In humans, ubiquitination systems contain 2 ubiquitin-activating enzymes (E1s), at least 37 ubiquitin-conjugating enzymes (E2s) and about 670 ubiquitin ligases (E3s), while deubiquitination systems contain about 100 deubiquitylating enzymes (DUBs)[1]. E3s specifically add ubiquitin to substrates and DUBs specifically remove ubiquitin from substrates, constituting the complex regulatory system of ubiquitination. Ubiquitinome is a high-throughput method for qualitative and quantitative identification of protein ubiquitination. Generally, the main enrichment strategies of ubiquitinated protein in ubiquitinome include using antibodies that directly recognize ubiquitin, using antibodies that recognize tags fused with ubiquitin and using di-Gly antibodies to recognize ubiquitinated peptides after trypsin digestion[2-5]. There has been a plethora of studies addressing that the uncontrollable expression and function of enzymes in ubiquitination system during aging destroy protein homeostasis and affect senescence related signaling pathways such as insulin and mTOR[6, 7]. In order to systematically study the changes of ubiquitination during aging, ubiquitinome has been applied to oncogene-induced senescence in human fibroblasts[8], ageing in C. elegans[9] and drosophila[10] in recent years.

Histone is an evolutionarily conserved DNA binding protein, whose family members H1, H2A, H2B, H3, and H4 can be ubiquitinated at multiple Lysine residues[11]. H2A is the earliest discovered ubiquitinated protein and can be ubiquitinated at K13, K15, K119, K125, K127, and K129, of which K119 has the highest frequency of ubiquitination[11, 12]. Many E3s and DUBs including RNF168, RNF8, BRCA1, USP3, USP51, and USP10 regulate the ubiquitination level of H2A[13, 14]. H2A ubiquitination mainly plays epigenetic regulation and DNA damage response (DDR) functions in cells[15]. For example, H2AK118/119ub plays a role in the initiation of DDR by inhibiting DNA transcription[16], while H2AK13/15ub can directly stabilize the DNA replication fork[17]. Due to the central role of DDR in aging and tumorigenesis, a majority of studies have focused on the function of H2A ubiquitination in aging and tumorigenesis. Through ubiquitinome, Yang et al. found that increased ubiquitination of H2A can serve as an evolutionarily conserved marker of aging[10]. Changes in H2A ubiquitination also occur in tumors. Experimental evidence indicates that H2AK119ub decreases in prostate cancer compared to normal tissue[18]. In addition, studies have attempted targeting increased H2AK119ub levels with PTC-596 to treat ovarian cancer[19].

However, there was no relevant report revealed the systematic changes of ubiquitination in mice during aging and the causes of such changes. In addition, Whether the ubiquitination changes during aging also exist in tumorigenesis and how they effect on tumor have remained unknown. Here, we obtained whole-life ubiquitinome data from mouse brain, heart, liver, muscle, and spleen, and systematically analyzed its relationship with RNA sequencing data. Furthermore, we comprehensively investigated the impact of H2A’ E3s/DUBs expression, which caused the greatest ubiquitination changes during aging in mice, on the prognosis of pan-cancer and its possible mechanism.

## Results

### Protein ubiquitination changes more intense during aging

To characterize and quantify the ubiquitination of proteins during aging in mice, a high-throughput method based on liquid chromatography-mass spectrometry (LC-MS) was used according to previous study[5]. Brain, heart, liver, muscle, and spleen of 1 month (M1), 6 month (M6), 12 month (M12), 18 month (M18), and 24 month (M24) mice were lysed and trypsinized to obtain short peptides. The short peptides containing ubiquitination modification site were enriched by specific antibodies and identified by LC-MS after tandem mass tag (TMT) labeling (Fig 1a). PCA analysis and correlation analysis showed that there were significant differences in ubiquitinome of different organs in mice, especially heart and liver (Fig 1b and Fig S1a). Among five organs, liver has the highest number of regulated ubiquitinated sites (fold change > 1.5) and spleen has the least number of regulated ubiquitinated sites (Fig 1c, Fig S1b and Table S1). In addition, compared with other organs, liver has a higher proportion of down-regulated ubiquitination sites than up-regulated ubiquitination sites and the brain has the opposite trend (Fig S1c). By using time fuzzy clustering method, regulated ubiquitinated sites could be classified into four groups: decreased in early aging; decreased in late aging; increased in early aging; increased in late aging (Fig 1d). As the mice grew older, the proportion of ubiquitinated sites with significantly regulated (fold change > 1.5) increased in almost all organs (Fig 1e and Table S1). Besides, the variation in ubiquitination levels of protein sites became more dramatic in all five organs with mice aging (Fig 1f and Fig S1d).

**Fig. 1.**
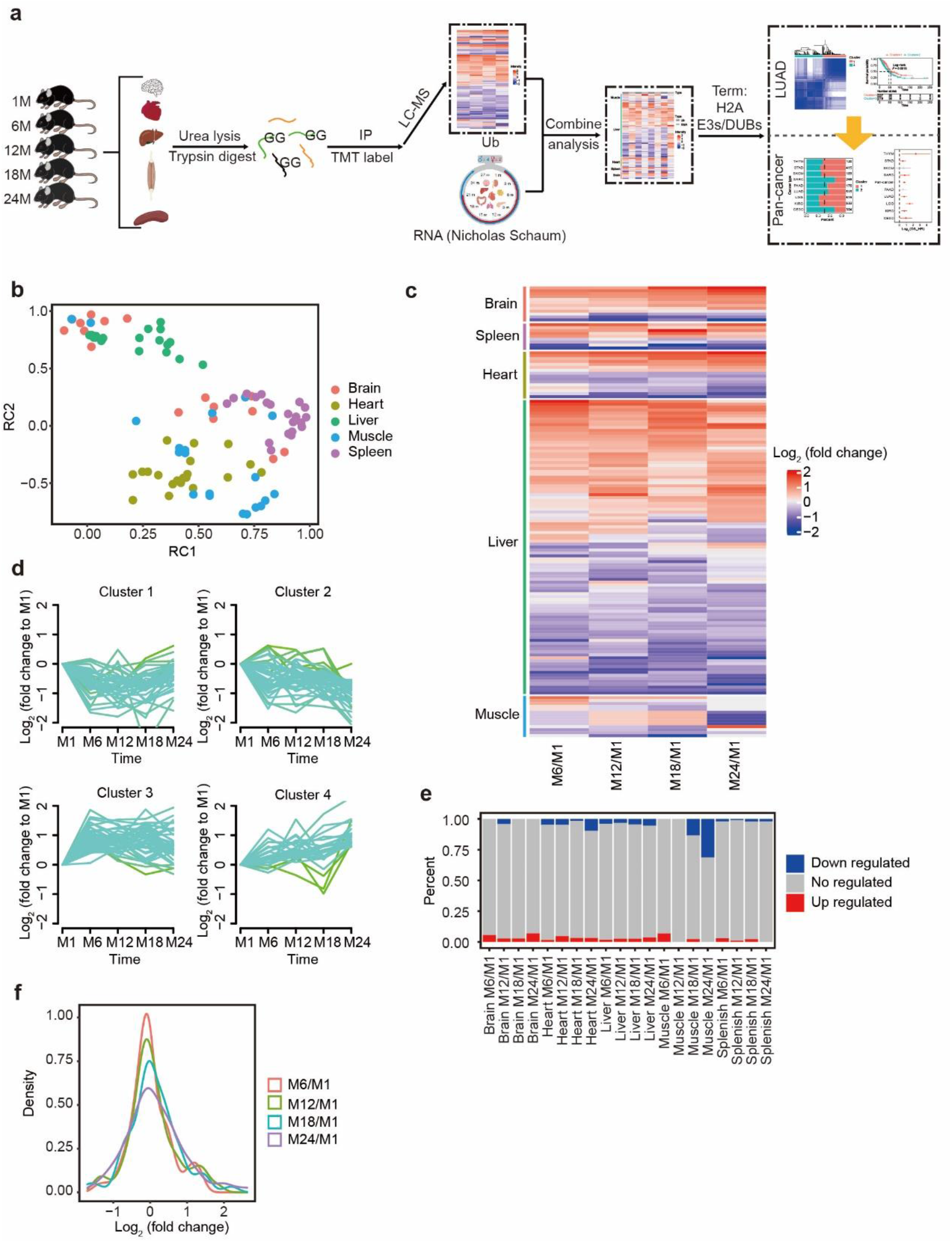
Ubiquitinome in mice aging. **#1** Study workflow. M month, GG glycine-glycine residues, TMT Tandem Mass Tag, LC-MS Liquid chromatography-mass spectrometry, Ub ubiquitinome, RNA RNA sequencing data, LUAD Lung adenocarcinoma. **b** Principal component analysis (PCA) of the ubiquitinome data from different mouse orangs. **c** Heatmap showing the mean of relative intensity of ubiquitinated sites in different ages to M1 in five mouse organs. **d** The regulated ubiquitinated sites are categorized to four clusters using fuzzy c-means algorithm, each line chart represents a pattern of ubiquitination modification changes with age. **e** Percentage of regulated and unregulated ubiquitinated sites in different mouse orangs and ages. One month (M1) is used as a reference age. **f** Density plots showing the distribution of the fold chang for intensity of ubiquitinated sites in mouse heart with aging.

### Protein ubiquitination changes during aging is caused by changes in expression of E3s/DUBs

To investigate the reasons of protein ubiquitination changes during aging, we integrated our ubiquitinome data with RNA sequencing data for all five organs and all five ages (Fig 1a). We found that RNA expression of proteins with significantly regulated ubiquitination during aging did not change (mostly in brain, heart, muscle and spleen) or the trend of change was opposite (mostly in liver), suggesting that the variation in ubiquitination was not due to gene expression (Fig 2a). Moreover, the correlation of protein ubiquitination changes and expression changes relative to M1 gradually decreased from 0.202 in M6, 0.152 in M12, 0.086 in M18 to 0.077 in M24 (Fig 2b). So, what exactly caused the changes in the ubiquitination of various organs in aging mice? We then analyzed the expression of four ubiquitin-expressing genes (Rps27a, Uba52, Ubb and Ubc) in mice and found their expression levels did not change with aging in all five organs, suggesting that the variations in ubiquitination were also not due to the expression of ubiquitin (Fig 2c). Some ubiquitination link types such as K48-link ubiquitination cause the degradation of proteins which can reduce the quantity of ubiquitinated protein, so we also analyzed the ubiquitination link type changes during mice aging. The result shows no differences in ubiquitination link type between distinct ages in all organs except heart (Fig S2a). After ruling out the above possible conjectures, we hypothesized that the expressions of enzymes in E3s/DUBs system probably affected the ubiquitination modification level of protein. We found that the variation trends of enzyme expression in E3s/DUBs system and ubiquitination levels were similar to each other than the total gene expression during mice aging (Fig 2d and Table S2). Besides, the transcription levels of enzymes in E3s/DUBs system were wildly fluctuated with age (Fig S2b and S2c).

**Fig. 2.**
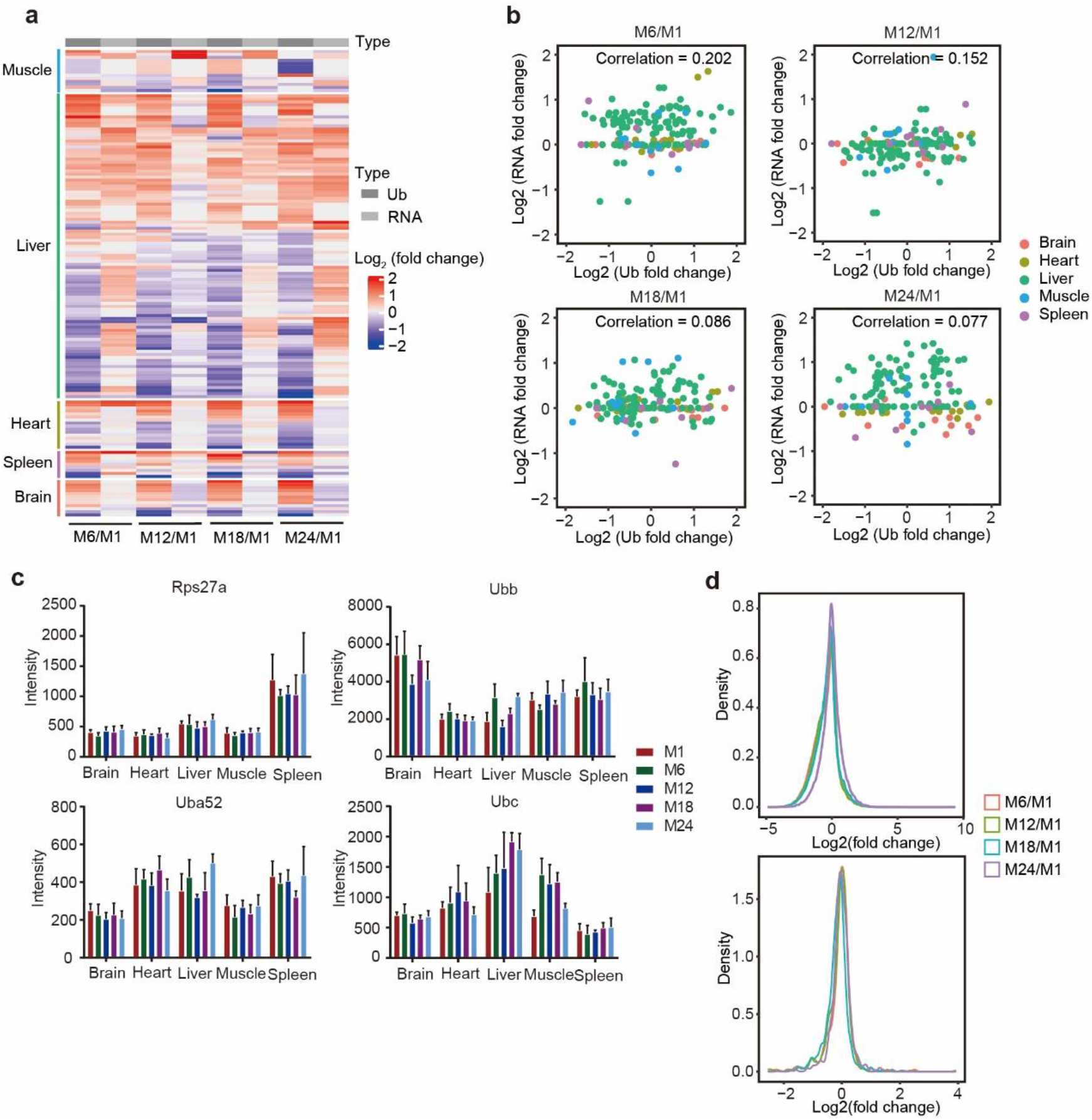
The changes of ubiquitinome in aging mice are not caused by gene expression. **#1** Heatmap shows the relationship between ubiquitination levels and gene expression levels for regulated ubiquitinated protein in various organs during aging in mice. each line represents the same protein. **b** Correlation plot of protein ubiquitination changes and gene expression changes in diverse age mouse. **c** Expression levels of four ubiquitin-coding genes in different age mouse. **d** Density plots showing the distribution of relative changes in total RNA (upper) and E3s/DUBs RNA (lower) during cardiac aging in mice.

### The ubiquitination of H2A was obviously regulated during aging

We then focused on which proteins were significantly altered in ubiquitination during aging in mice. Firstly, GO (cell components, biology process and molecule function) and KEGG enrichment analyses were performed for proteins with significant ubiquitination changes. The results show nucleosome, DNA packaging complex and protein-DNA complex terms were enriched (Fig 3a and S3a-c). Besides, GSEA enrichment analyses also found chromatin organization, nuclear chromosome, nuclear chromatin and chromosome organization terms were enriched (Fig 3b and S3d). The results of both enrichment analyses suggested that the ubiquitination of histone was significantly altered during aging. The line diagram shows that a significant proportion of the ubiquitination intensity of histones varies with aging in the five organs (Fig 3c and S3e). By analyzing the alteration of ubiquitination modification of histone in M24 relative to M1, we found four of the six histones (H2ax, H2ac12, H2ac20, H2aj, H2bc7 and H3c8) with significantly regulation belong to H2A (Fig 3d). As expected, the transcription level of enzymes in E3s/DUBs of H2A were wildly fluctuated during aging in mice, which was opposite to the change trend of transcription level of total genes (Fig 3e, 3f and Table S3). In light of the results obtained above, the transcription of E3s/DUBs of H2A significantly altered with mice age, leading to regulated ubiquitination of H2A.

**Fig. 3.**
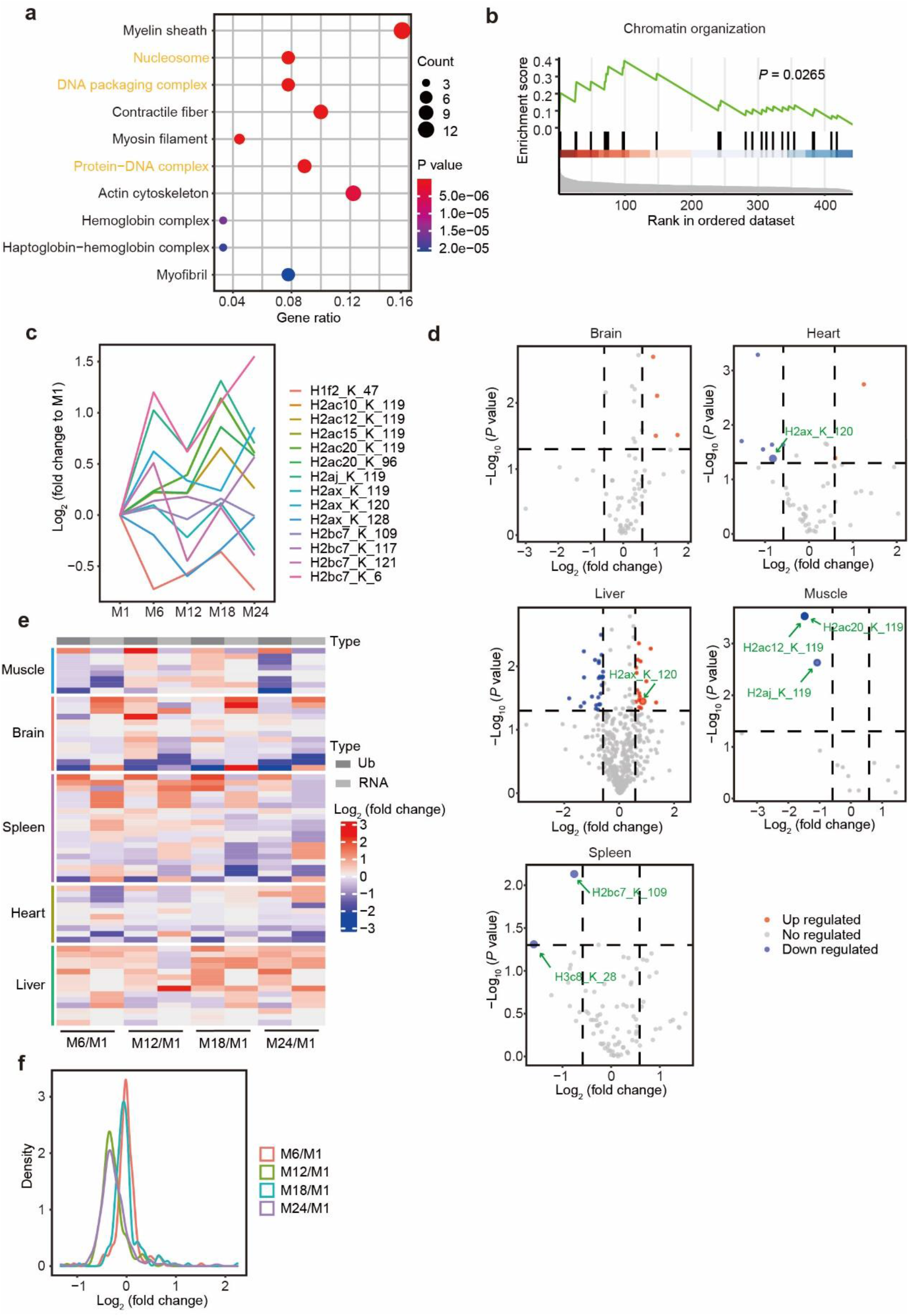
The ubiquitination of H2A in aged mice was significantly changed. **#1** Bubble chart showing the result of GO cell components enrichment analysis of proteins with regulated ubiquitination during mice aging. **b** GSEA analysis result for ubiquitinome from mouse liver (M24 vs M1). **c** Relative abundance of histone ubiquitination levels in liver of mice in different age (relative to M1). **D** Volcanic map shows proteins with significant ubiquitination changes in M24 relative to M1 in five mouse organs. **e** Heatmap shows the relationship between ubiquitination levels and gene expression levels for histone in various organs during aging in mice. each line represents the same protein. **f** Density plots showing the distribution of relative changes in gene expression of H2A’s E3s/DUBs during brain aging in mice.

### Construct transcriptomic subtypes by the E3s/DUBs of H2A in LUAD

The occurrence and development of tumor is closely related to aging. In the previous analysis, we found that the alteration of the expression of H2A’s E3s/DUBs was the key factor leading to the alteration of ubiquitination during aging. But what role does H2A’s E3s/DUBs play in tumors? We then analyzed bulk RNA-sequencing data from 506 LUAD patients in TCGA. Utilizing unsupervised consensus clustering of gene expression values of H2A’s E3s/DUBs, we identified two transcriptomic classes 1-2 (Fig 4a and Table S4). Kaplan-Meier survival analysis showed significantly different overall survival (p =0.0018; log-rank test) between the two classes: patients with class 1 tumors displayed good outcome, whereas patients with class 2 tumors had poor prognosis (Fig 4b). Transcriptomic classes were not significantly associated with sex and age (Fig 4c). In the analysis of gene set variation analysis (GSVA) of human hallmark gene sets and curated KEGG gene sets from GSEA official website, we discovered that class 2 tumors highly expressed genes participating in nucleotide excision repair, DNA repair, base excision repair, DNA replication, mismatch repair and homologous recombination (Fig 4c). The high expression of genes related to DNA repair suggests that the genome of class 2 tumors may be instability, which was consistent with the short overall survival time. We also estimated the presence of immune cells in tumor microenvironment by xCell method. The results show class 2 tumors had significantly more Th1 and Th2 cells compared to class 1 (Fig 4c).

**Fig. 4.**
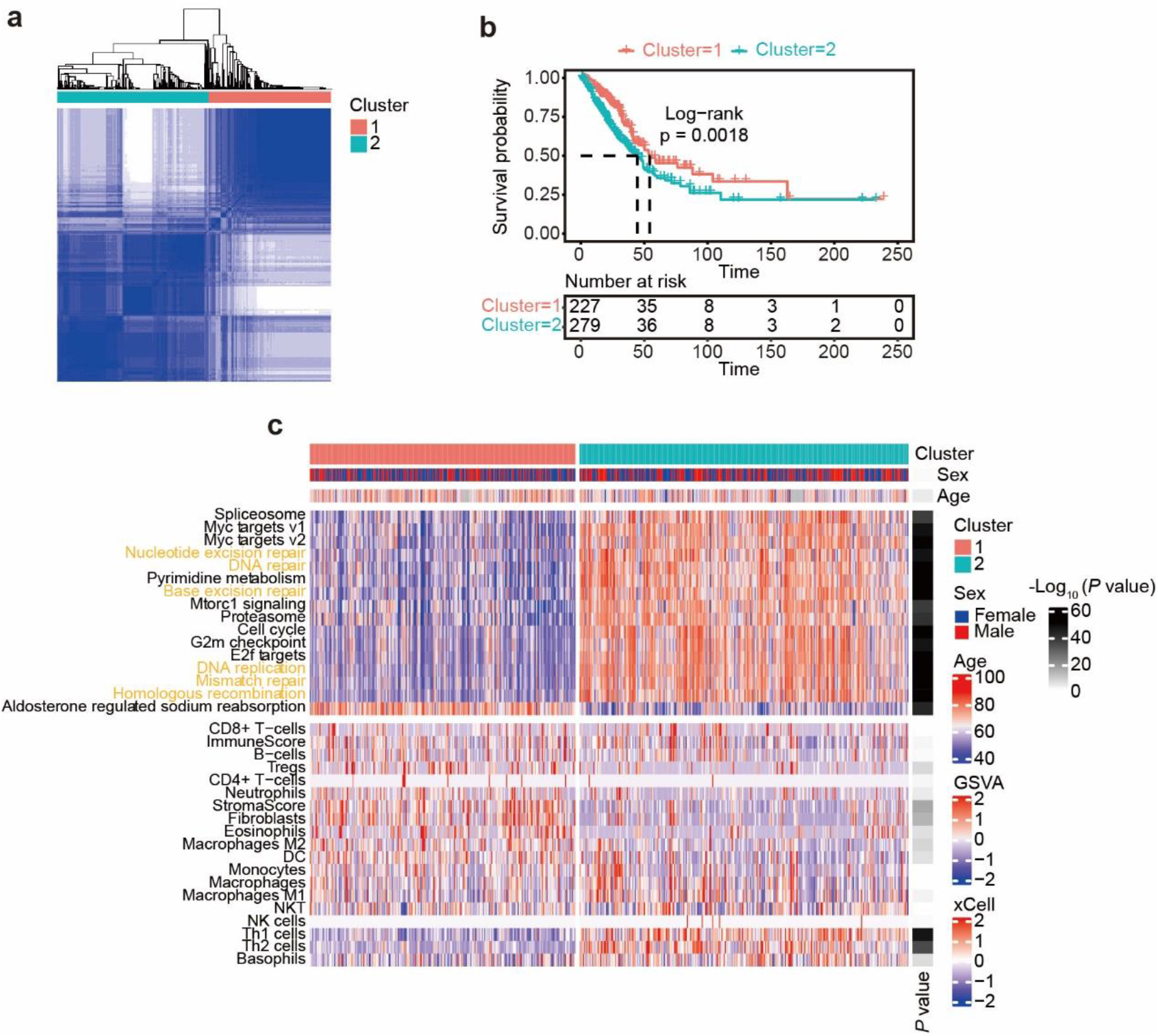
E3s/DUBs of H2A are applied to transcriptomic class in LUAD. **#1** Consensus matrix for two clusters of 506 LUAD patients stratified by the expression of H2A’s E3s/DUBs. White color represent two sample never clustered in one cluster and blue color represent two sample always clustered in one cluster. **b** Kaplan–Meier plot of overall survival for LUAD patients stratified by transcriptomic class 1–2. **c** GSVA (The 16 terms with the greatest differences between classes 1–2 are displayed) and immune infiltration analysis for transcriptomic class 1–2.

### Significant differences in genomic stability between transcriptomic subtypes

In previous analyses, we have found that class 2 tumors have higher expression of DNA repair related gene than class 1 tumors. Therefore, we wanted to further clarify whether this difference is associated with genomic stability. Genome-wide mutation landscapes of LUAD reveal that mutations including nonsense mutations, missense mutations, splice mutations, frame mutations and synonymous mutations were all much more in class 2 tumors than class 1 (Fig 5a). Class 2 tumors were distinctly associated with high mutation of oncogenes like TP53 and IRS1 (Fig 5a)[20]. Besides, Genome-wide copy number landscape of LUAD disclose both gain and loss copy number alteration were more evident in class 2 tumors than class 1 tumors (Fig 5b). The most obvious differences are copy number gain on chromosomes 2, 3, 7, 8 and 17, and copy number loss on chromosomes 5 and 16 (Fig 5b). From the perspective of gene level, tumor suppressor RASA1 exhibited less gain copy number in class 2 tumors than class 1 tumors (Fig S4a). Epigenetic analysis showed that tumor suppressor SETD2 had a high methylation level in class 2 tumors, which inhibited its gene expression and resulted in reduced tumor suppressive function (Fig 5c). Throughout the full genome-wide landscape of methylation, difference between class 1 tumors and class 2 tumors is not evidential (Fig S4b).

**Fig. 5.**
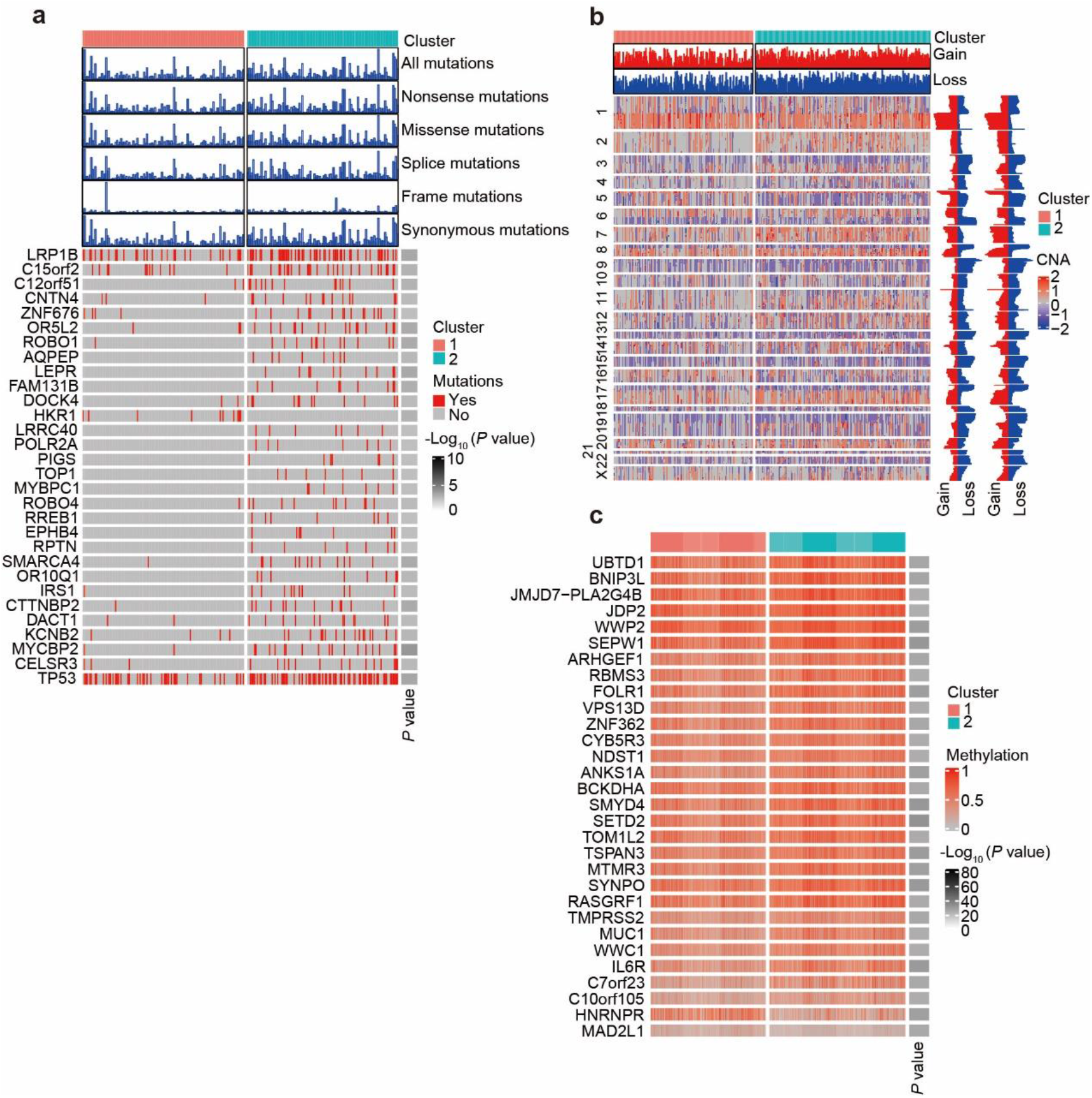
Mutation, copy number and methylation discrepancy in LUAD subtypes. **#1** Genome-wide mutation landscape of LUAD stratified by transcriptomic class 1–2. Top 30 genes with significantly different ratio in mutation between two classes are list. **b** Genome-wide copy number landscape of LUAD stratified by transcriptomic class 1–2. Gain and loss for class 1 (left) and class 2 (right) are summarized on the right panel. Genes are arranged in the order they are located on the chromosome (show on left of heatmap). **c** Heatmap of top 30 genes with significantly different methylation levels between transcriptomic class 1–2.

### The two transcriptomic subtypes are conserved in pan-cancer

We analyzed transcriptomic data from additional eight cancer types to determine whether the identified two transcriptomic subtypes also exist in other cancers. After unsupervised consensus clustering of gene expression values of H2A’s E3s/DUBs, eight cancer types were also clustered to two transcriptomic classes 1-2 (Fig S4c and Table S4). In pan-cancer, transcriptomic class 1 was slightly more than transcriptomic class 2 with LGG having the most percent of transcriptomic class 1 and SARC having the least percent of transcriptomic class 1 (Fig 6a). Compared with transcriptomic cluster 1, transcriptomic cluster 2 had significantly worse overall survival in 7 of 9 cancers (*P* < 0.05, cox proportional-hazards model) (Fig. 6b) and worse disease free survival in 4 of 9 cancers (*P* < 0.05, cox proportional-hazards model) (Fig. S4d). We then did the Kaplan-Meier survival analysis of overall nine cancer types, the results show patients with transcriptomic cluster 1 tumors had better outcome than patients with transcriptomic cluster 2 tumors for both overall survival (Fig. 6c) and disease free survival (Fig. S4e). Importantly, we found GSVA and immune infiltration analysis results in pan-cancer were the same as in LUAD. Specifically, the class 2 tumors were more active in terms including nucleotide excision repair, DNA repair, base excision repair, DNA replication, mismatch repair and homologous recombination (Fig 4c). In addition, like LUAD, the infiltration score of Th2 cells which promoted tumor immune evasion and metastasis in class 2 tumors were also dramatically higher than class 1 tumors in pan-cancer (Fig 4c) [21-23].

**Fig. 6.**
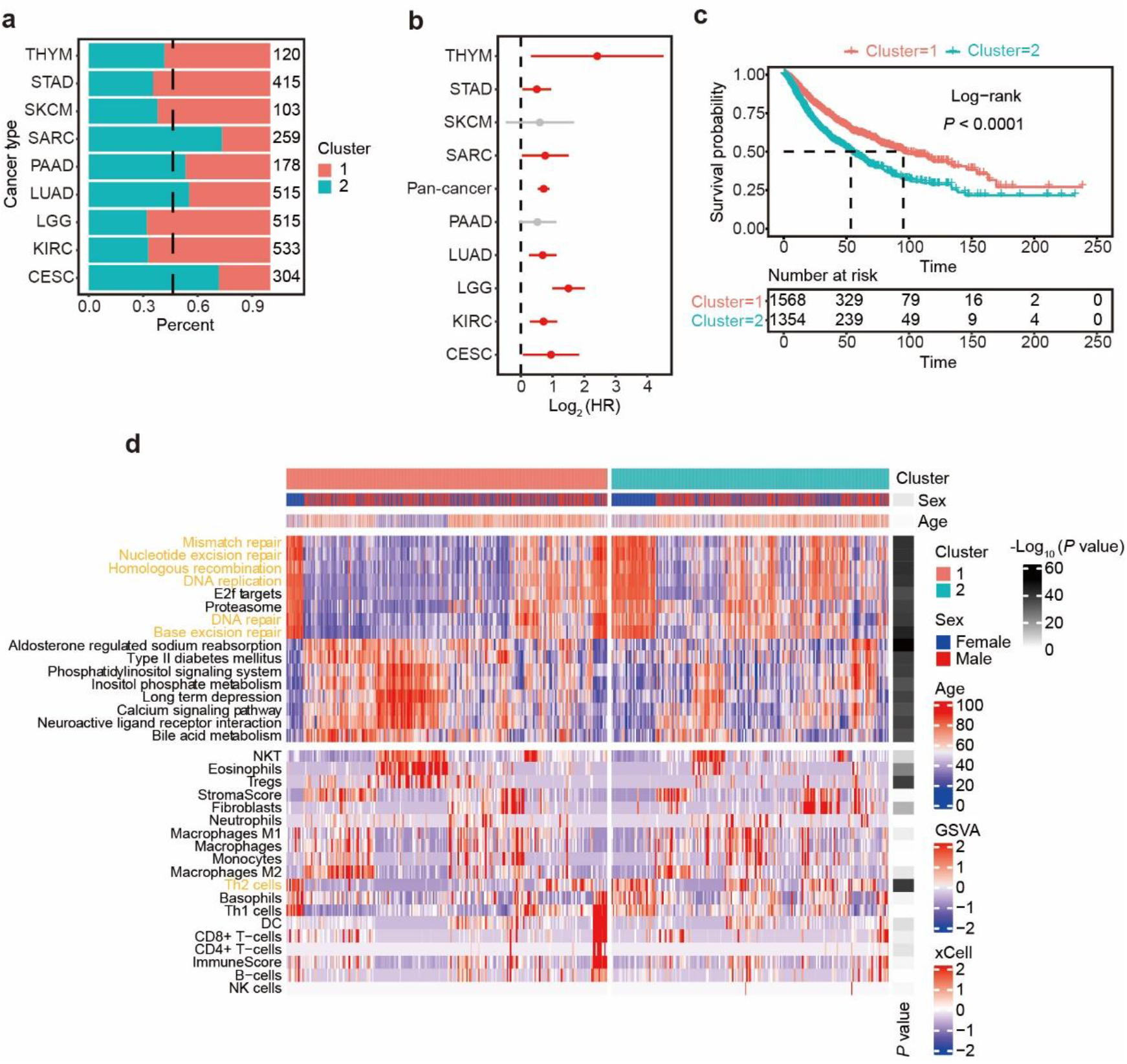
E3s/DUBs of H2A are applied to transcriptomic class in pan-cancer. **#1** Bar chart shows the patient number of class 1 and class 2 of the nine types of cancer after transcriptomic class. **b** Associations of transcriptomic subtypes and pan-cancer overall survival. The hazard ratios (HR) are calculated by the trend association (clusters 1–2) in cox proportional hazard models. **c** Kaplan– Meier plot of overall survival for pan-cancer patients stratified by transcriptomic class 1–2. **d** GSVA and immune infiltration analysis for transcriptomic class 1– 2.

## Discussions

Our study provides temporal resolution of ubiquitinome for five important organs across the overall lifespan of mice, providing fundamental resource for aging related research. Notably, we recognized that the intense changes happened in protein ubiquitination during aging might have little bearing on the protein expression itself, but on related E3s/DUBs. Furthermore, we demonstrated that the ubiquitination of H2A changed most obviously during aging, and the E3s/DUBs of H2A played an important role in pan-cancer outcomes. Our systemic evaluation of transcriptome, genome-wide mutation landscape, genome-wide copy number landscape and genome-wide landscape of methylation revealed the role of E3s and DUBs of H2A in influencing tumor prognosis by altering genomic stability. The unstable genome leads to a high tumor mutational burden, a high infiltration of tumor immune escape associated cells like Th2 cells, and therefore a high sensitivity to immunotherapy[24, 25]. Compared with the previous RNA sequencing database of aging mouse[26], one of the shortcomings of this study is that only five organs were utilized. Therefore, ubiquitinome of other mouse organs during aging should be considered in the future. In this study, RNA sequencing data rather than proteome data were used to represent protein expression, which needs to be improved in subsequent studies because the post-transcriptional regulation of proteins was ignored. In addition, although the expression of E3s and DUBs of H2A was significantly changed during aging and was used for classification of pan-cancer based on the fact that aging is an important risk factor for tumor, there was no apparent difference in age between the two tumor classes[27]. We speculate that the reason might be that the “senescence” of tumor cells fails to synchronize with the aging of human.

In summary, our study indicated the indispensable role played by H2A ubiquitination and H2A associated E3s/DUBs in aging and tumor progression, opening up new avenues for potential applications in the development of biomarkers for tumor prognosis and measurement of immunotherapeutic response.

## Figures and figure legends

**Fig. S1.**
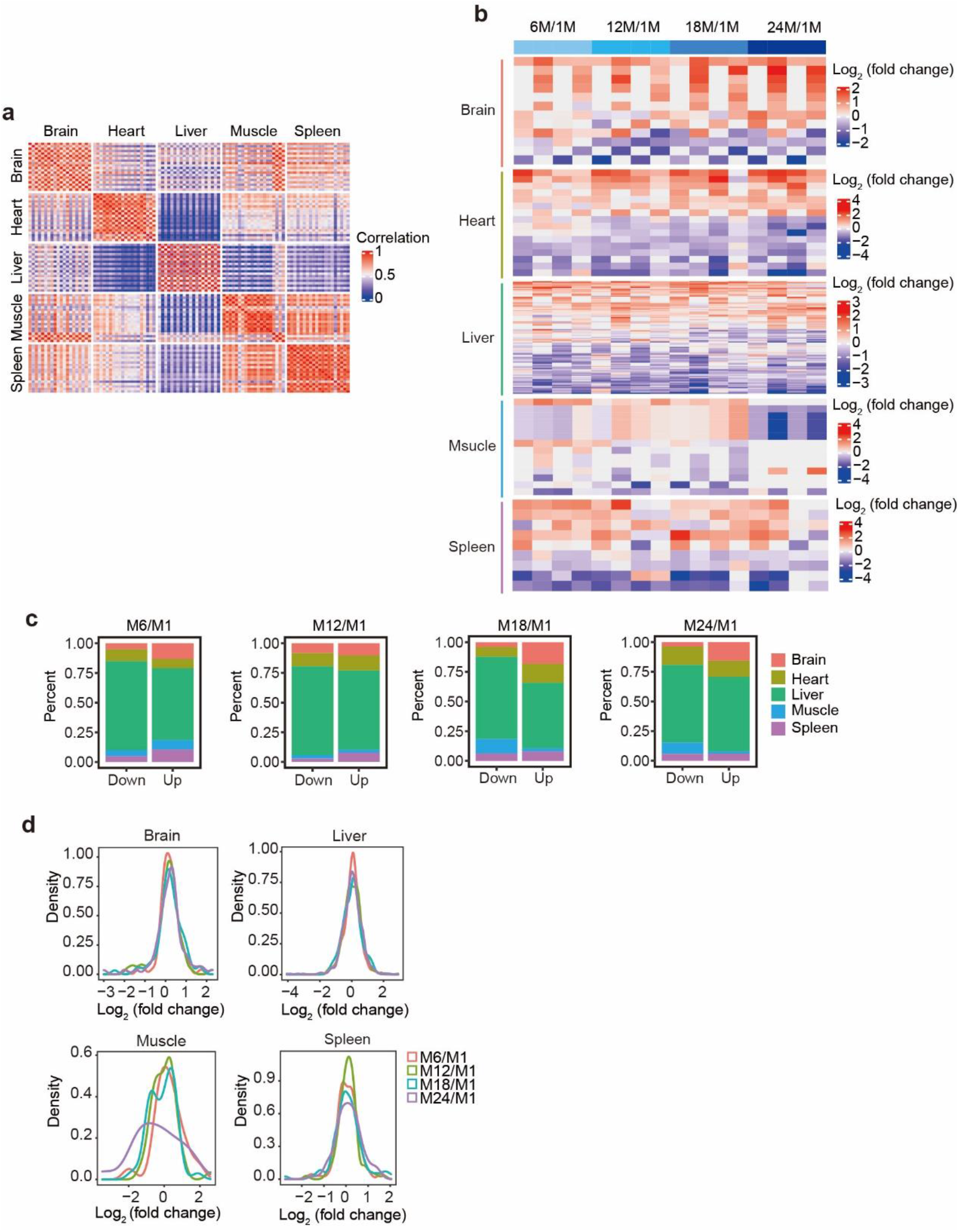
Ubiquitination omics of multiorgan and multiage mice. **a** correlation plot of ubiquitinome data from five mice organs. **b** Heatmap showing the relative intensity of ubiquitinated sites in different ages to M1 in five mouse organs. **c** Histogram shows the organ distribution of regulated ubiquitinated sites in mice of different ages. **d** Density plots showing the distribution of the fold change for intensity of ubiquitinated sites in mouse brain, liver, muscle and spleen with aging.

**Fig. S2.**
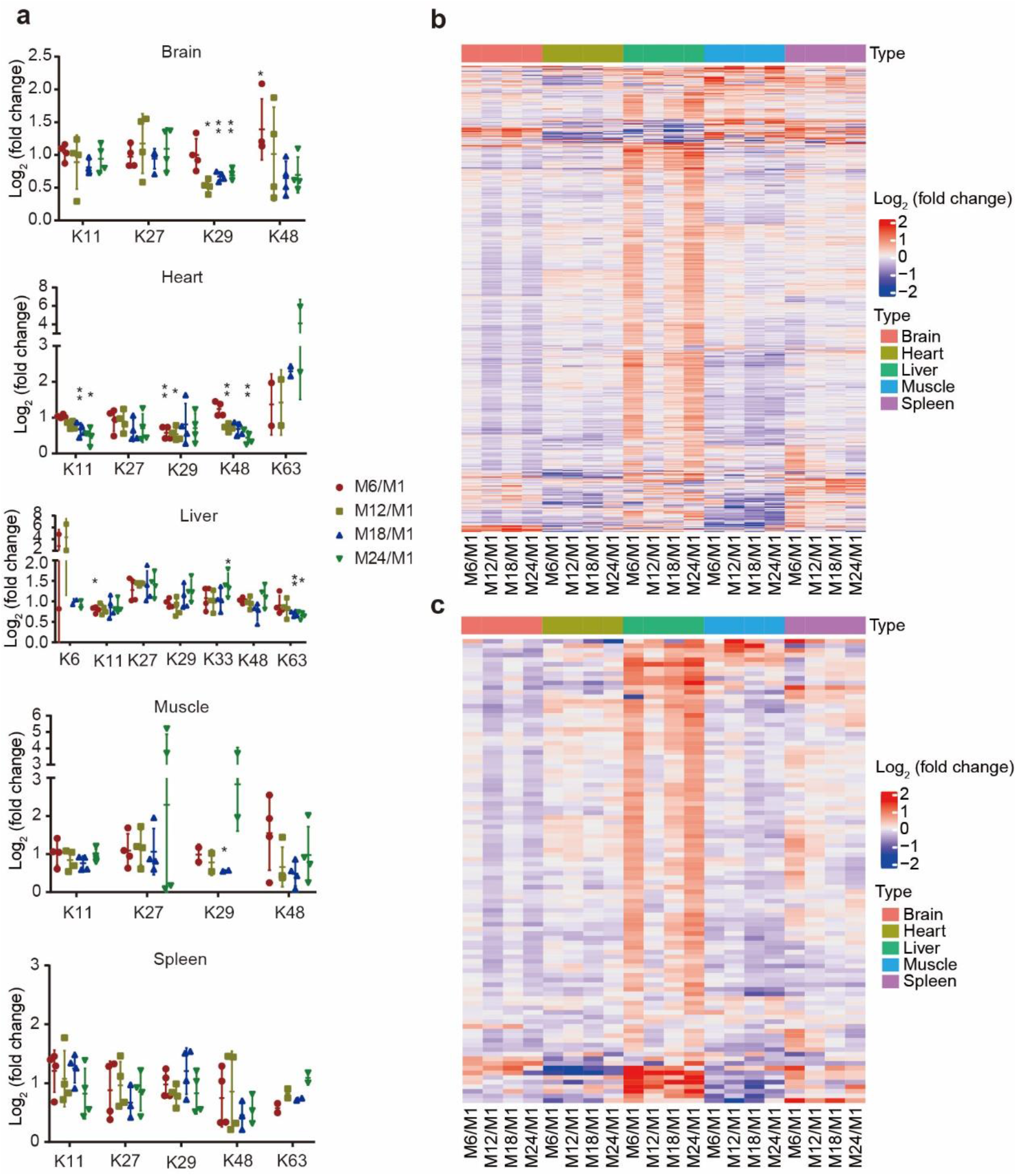
Integrated analysis of ubiquitinome and transcriptome during aging in mice. **a** Dot plot showing the intensity of seven ubiquitin chain linking patterns change with age in five mouse organs. **b-c** Heatmap of the transcription level of E3s/DUBs across different mice orangs and ages.

**Fig. S3.**
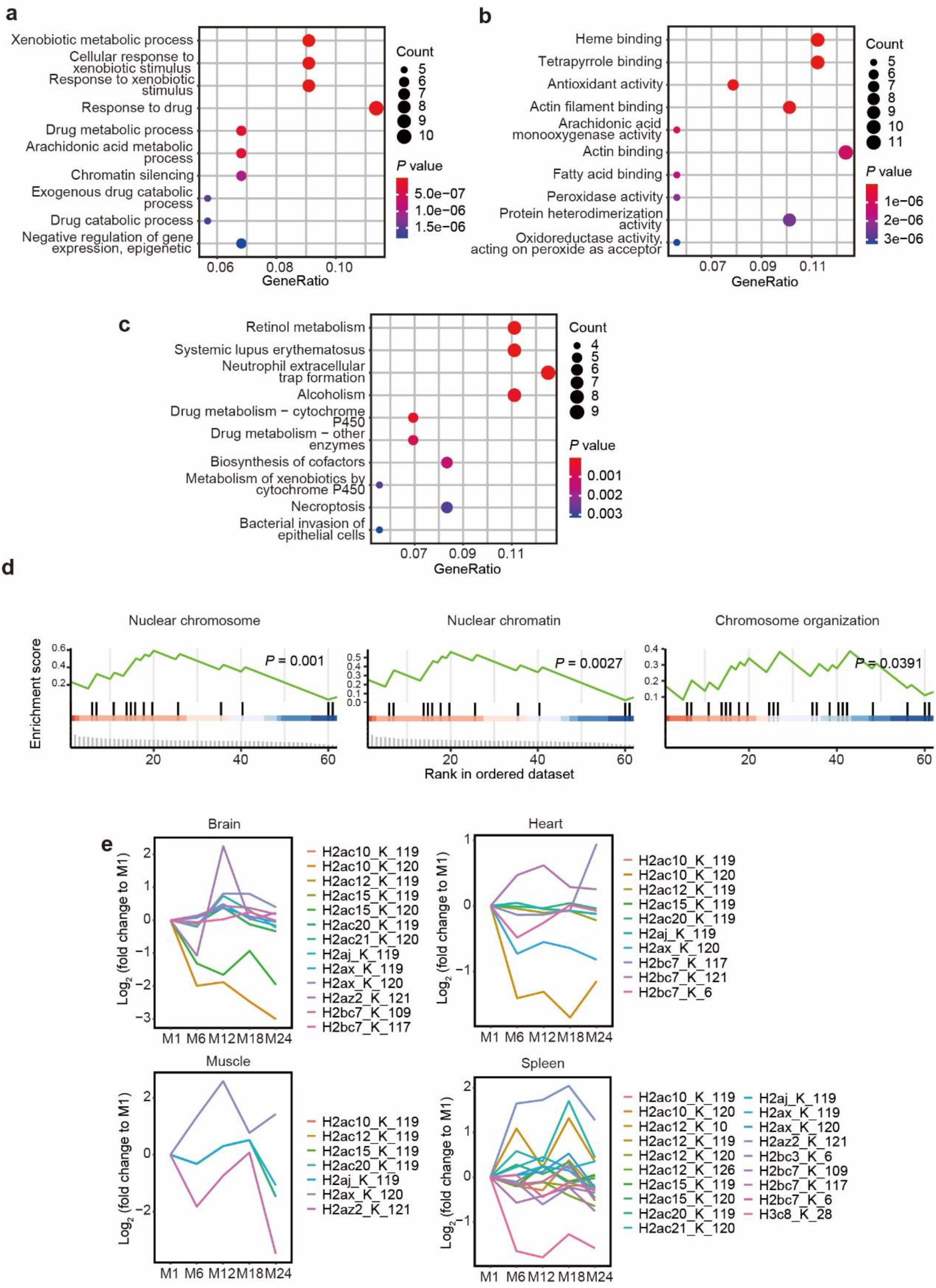
Enrichment analysis of ubiquitinome. **a-c** Bubble chart showing the result of GO biology process, GO molecule function and KEGG enrichment analysis of proteins with regulated ubiquitination during mice aging. **d** GSEA analysis result for ubiquitinome from mouse brain (M12 vs M1). **e** Relative abundance of histone ubiquitination levels in brain, heart, muscle and spleen of mice in different age (relative to M1).

**Fig. S4.**
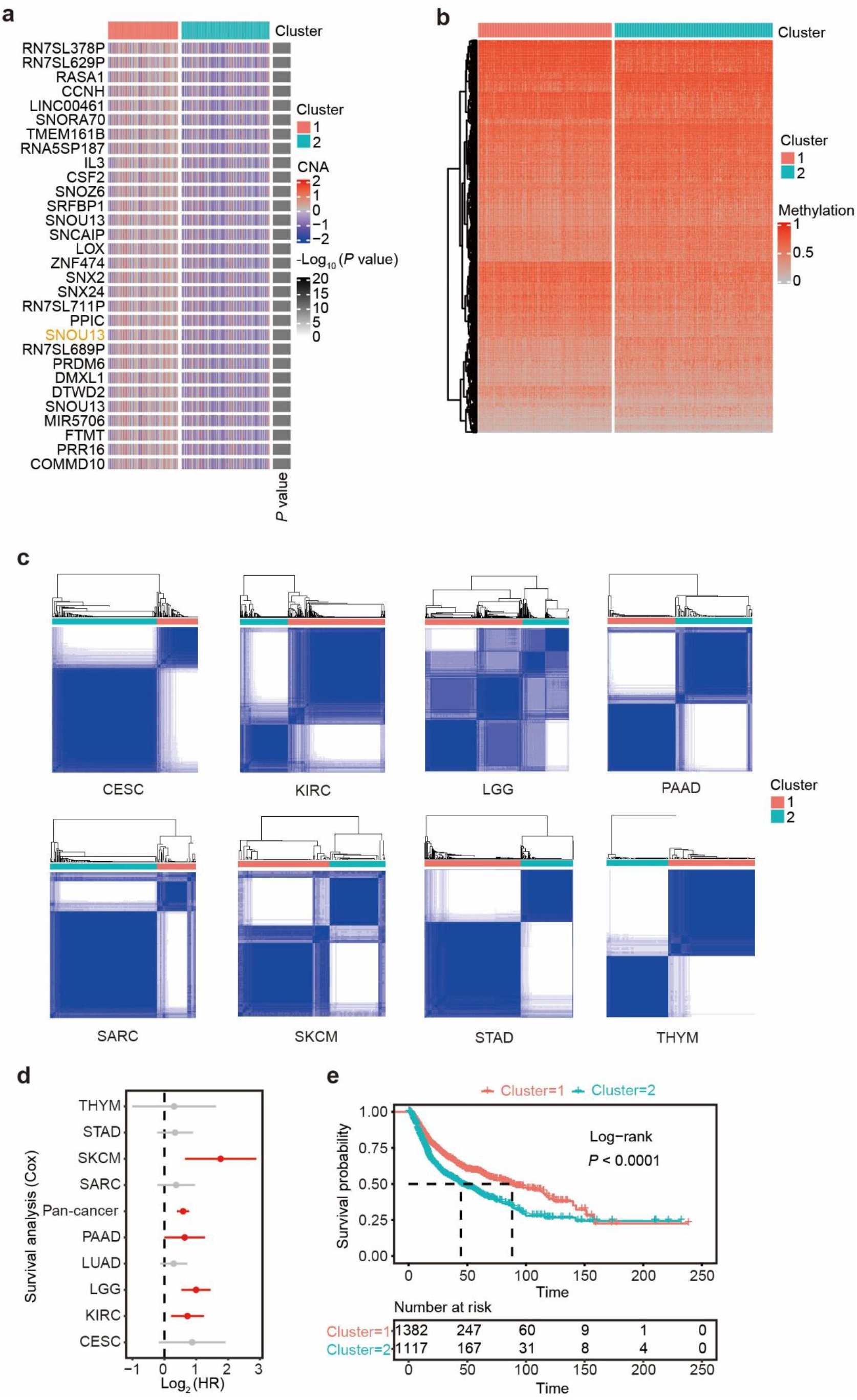
Transcriptomic class utilize E3s/DUBs of H2A. **a** Heatmap of top 30 genes with significantly altered copy number between transcriptomic class 1–2 in LUAD. **b** Heatmap showing genome-wide discrepancy of methylation between transcriptomic class 1–2 in LUAD. **c** Consensus matrix for two clusters of eight cancer types stratified by the expression of H2A’s E3s/DUBs. White color represent two sample never clustered in one cluster and blue color represent two sample always clustered in one cluster. **d** Associations of transcriptomic subtypes and pan-cancer disease free survival. The hazard ratios (HR) are calculated by the trend association (clusters 1–2) in cox proportional hazard models. **e** Kaplan–Meier plot of disease free survival for pan-cancer patients stratified by transcriptomic class 1–2.

## Materials and methods

### Reagents

Protease inhibitor (Cat. P1049) was purchased from Beyotime Biotechnology (Shanghai, China). Trypsin (Cat. T1426), urea (Cat. 5378), dithiothreitol (Cat. 43816), Iodoacetamide (Cat. A3221) and trifluoroacetic acid (TFA, Cat. 302031) were obtained from Sigma-Aldrich (Darmstadt, Germany). Tris-HCl (Cat. B548127), sodium chloride (Cat. A610476) and ammonium bicarbonate (Cat. A610032) were provided by Sangon Biotech (Shanghai, China). Besides, PR-619 (Selleck Chemicals, Cat. S7130, USA), PTMScan Ubiquitin Remnant Motif Kit (Cell Signaling Technology, Cat. 5562, USA), C18 Sep-Pak SPE cartridge (Waters, Cat. WAT023590, USA) and TMT Label Reagents (Thermo Scientific, Cat. 90110, USA) were used in our study.

### Animals

The Animal Care Committee of the Animal Experimental Ethical Inspection of the First Affiliated Hospital, College of Medicine, Zhejiang University approved the animal experiment protocols (permit number: 2018-841). Male C57BL/6 mice were obtained from the Experimental Animal Center of Zhejiang Academy of Medical Sciences. Food and water were freely provided to the mice. They were housed in the Laboratory of Animal Center of the First Affiliated Hospital, College of Medicine, Zhejiang University, at standard conditions in a 12-hour photoperiod. Mice were euthanized and five tissues including liver, brain, heart, skeletal muscle and spleen were collected at 1, 6, 12, 18, 24-month-old. This study was reviewed and approved by the tab of Animal Care Committee of the Animal Experimental Ethical Inspection of the First Affiliated Hospital, College of Medicine, Zhejiang University (permit number: 2018-841).

### Protein extraction and processing

About 100 mg tissues were lysed by freshly prepared 4 °C lysis buffer (8 M urea with 150 mM NaCl, 50 mM pH 8.0 Tris-HCl, protease inhibitor and PR619). After centrifuge at 15000 × g for 15 min at 4 °C, the supernatant was reduced by dithiothreitol and stopped by iodoacetamide. Besides, the protein concentration was calculated by the bicinchoninic acid protein assay (Biosharp, China). About 5 - 10 mg protein was used. The protein was diluted with PBS to 2 µg/µL, adding 5 times the volume of -20 °C pre-cooled acetone and incubate at -20 °C for at least 4h. After centrifuging at 3000 × g for 10 min, remove the supernatant and resuspend in 100mM ammonium bicarbonate buffer with ultrasonic solubilization. Then the protein samples were digested overnight at 37 °C with a trypsin-to-substrate ratio of 1:40 (wt/wt) and shaken at 150 rpm. After digestion, peptides were acidified with trifluoroacetic acid (TFA) to a final concentration of 1%. Acidified peptides were centrifuged at 1800 × g for 10 min at 4 °C to remove lipids, and the supernatant was desalted by using C18 Sep-Pak SPE cartridge (Waters, USA) according to the manufacturer’s instruction. The eluate buffers were dried by vacuum centrifugation.

### Enrichment of ubiquitinated peptides

The peptides were resuspended in cold 1 × IAP buffer and incubated with ubiquitinated cross-linked antibodies for 2 h at 4 °C according to our pervious experiments [5]. Then, the IP tube was centrifuged at 100 × g for 30 s at 4 °C to gather beads which were washed thrice with cold 1 × IAP buffer and twice with cold mass spectrum (MS) water. diGly-modified peptides were eluted by adding 50 μL 0.15% TFA twice and dried by vacuum centrifugation.

### Tandem mass tag (TMT) isobaric labeling

Equilibrate the TMT labeling reagents to room temperature and add 84 μL acetonitrile to dissolve the reagent within 5 min by vortexing occasionally. diGly-modified peptide sample was dissolved in 20 μL 100 mmol/L tetraethylammonium bromide (TEAB, pH = 8.0) and 8 μL TMT label reagent was added separately. The mixed buffer was incubated for 1 h at room temperature. Then, 2 μL 5% hydroxylamine was added to each sample and incubated for 15 min to stop the reaction. The supernatant was desalted by a homemade C18 stage tip and dried by vacuum centrifugation.

### Liquid chromatography-tandem mass spectrometry (LC-MS) analysis

TMT labeled peptide was dissolved in solvent A (0.1% FA in H_2_O), loaded onto an Acclaim PepMap 100 C18 trap column (Dionex, 75um×2cm) by Ultimate 3000 nanoUPLC (Dionex) and eluted onto an Acclaim PepMap RSLC C18 analytical column (Dionex, 75um×25cm). The gradient of solvent B (0.1% FA in 80% ACN) was 3% in 0-4 min, 3% to 5% in 4-6 min, 5% to 15% in 6-70 min, 15% to 30% in 70-90 min, increased to 80% in 10 min and then keep 80% for 10 min, decreased to 3% in 10 min finally, the flow rate was 300 nl/min. The peptide was analyzed by Q Exactive HFX (Thermo Scientific) with resolution of 70000, the m/z scan range of 350 to 1500, a data-dependent top 20 precursor ions, a threshold of 5E4 with 15s dynamic exclusion. Peptide was fragmented using a collision energy of approximately 32% normalized collisional energy; ion fragments were detected at a resolution of 45000; 3E6 ions were accumulated for generation of MS spectra and 1E5 ions for generation of MS/MS spectra.

### diGly-modified peptides identification and quantification

We utilized MaxQuant (version 2.0.3.0, https://www.maxquant.org) for diGly-modified peptide identification and quantification. The mouse UniProtKB database (October 2021) was used as search database. oxidation (M) (+15.99491 Da), acetylation (protein N-term) (+42.01056 Da) and GlyGly(K)_10plex_TMT (+343.20586 Da) was set as the variable modification, and Carbamidomethyl (C) (+57.02146 Da) was set as the fixed modification. The maximum number of missed cleavage sites in one peptide was set to 2. Digestion enzyme was trypsin with maximum number of 2 missed cleavage sites.

### Enrichment analysis

We performed enrichment analysis using R package clusterProfiler[28]. Proteins containing regulated ubiquitination were selected for cell component, biology process, molecule function, and KEGG enrichment analysis. The top 10 terms with the most obvious differences in each analysis were selected according to the P values, and arranged according to the P values from small to large. GSEA (using the R package clusterProfiler) analysis was used to confirm the reliability of the above pathway enrichment analysis. Besides, transcriptome data was used to performed geneset variation analyses using the R package GSVA [29] to obtain pathway enrichment scores for each tumor sample.

### Consensus clustering

The transcriptome data from CESC, KIRC, LGG, LUAD, PAAD, SARC, SKCM, STAD, and THYM was filtered to only include E3/DUBs of H2A. Consensus clustering was performed for each cancer types using the R package ConsensusClusterPlus [30] with parameters: maxK = 4, reps = 1000, pItem = 0.95.

### Immune infiltration analysis

The tumor immune cell infiltration was evaluated for CESC, KIRC, LGG, LUAD, PAAD, SARC, SKCM, STAD, and THYM using R package xCell [31]. The result was filtered to only include 19 immune-related cells and scores.

### Bioinformatics and statistical analysis

All computations and analyses were implemented in R 4.1.0 and relevant packages. Briefly, principal component analysis was performed using psych package. Fuzzy c-means clustering was performed using Mfuzz package. The pearson’s correlation was calculated using cor method in base package. The cox regression and log-rank test were performed using survival package. The chi-square test, Mann–Whitney–Wilcoxon test and t–test was selected to hypothesis testing for categorical and continuous data. *P* or BH adjusted *P* < 0.05 was regard as statistically significant.

## Acknowledgements

We thank Linbo Wang and Jichun Zhou for helpful discussions. This work was supported by National Key Research and Development Program of China (2021YFC2301805), National Natural Science Foundation of China (82103499).

## Authors’ contributions

FZ, ZW and JZ performed the majority of experiments. FJ, FZ and MZ performed the majority of data and statistical analysis. FJ and ZS directed the study. YC and SZ collected aging mice tissues.

## Availability of data and materials

The ubiquitinome data have been deposited to the ProteomeXchange Consortium (http://proteomecentral.proteomexchange.org) via the iProX partner repository[32] with the identifiers PXD035091. RNA sequencing data from various organs of mice of different ages were obtained from previous study[26]. The transcriptome, gene mutation, copy number alteration, and methylation data of CESC, KIRC, LGG, LUAD, PAAD, SARC, SKCM, STAD, and THYM were obtained from TCGA Firehose via cBioPortal[33, 34].

## Declarations

### Ethical approval and consent to participate

Not applicable.

### Consent for publication

All authors have reviewed and approved this manuscript.

